# Functional impact of pathogenic Runt domain mutations in *Runx2* on skeletal and dental development in cleidocranial dysplasia

**DOI:** 10.1101/2025.06.18.660258

**Authors:** Saki Ogawa, Shinnosuke Higuchi, Yuki Yoshimoto, Mari Hoshino, Shigenori Miura, Atsuko Hamada, Hitomi Watanabe, Tetsushi Sakuma, Kadi Hu, Shun Ogata, Kenta Uchibe, Katsumi Fujimoto, Takashi Yamamoto, Tetsuji Okamoto, Ryo Kunimatsu, Yusuke Sotomaru, Kotaro Tanimoto, Gen Kondoh, Toshihisa Komori, Denitsa Docheva, Chisa Shukunami

## Abstract

Runt-related transcription factor 2 (RUNX2) is essential for skeletogenesis, and mutations in its gene cause cleidocranial dysplasia (CCD), an autosomal dominant skeletal disorder. The evolutionarily conserved 128-amino acid Runt homology domain (RHD) of human RUNX2 is essential for DNA binding and heterodimerization, and serves as a mutation hotspot associated with severe CCD phenotypes. To elucidate the functional impact of pathogenic RHD mutations *in vivo*, we generated two novel mouse lines: one carrying a missense mutation, c.695G>A (p.R232Q) (*Runx2^m/+^*), corresponding to the human *RUNX2* c.674G>A (p.R225Q), and the other harboring a frameshift mutation, c.697_698delGA (p.E233TfsTer9) (*Runx2^112/+^*), causing a premature stop codon. Homozygous *Runx2^m/m^*and *Runx2^112/112^* mice lacked membranous ossification, whereas heterozygous *Runx2^m/+^* and *Runx2^112/+^* mice displayed typical CCD-like skeletal features, including an open anterior fontanelle and clavicle hypoplasia. Unexpectedly, heterozygotes carrying pathogenic mutations in RHD developed an accessory root-like protrusion at the furcation of three-rooted maxillary first molars, representing a previously unrecognized dental phenotype during root development. Dual luciferase assays revealed impaired transactivation of the p.R232Q mutant Runx2 on the osteocalcin enhancer/promoter. Wild-type Runx2 was robustly expressed in osteoblasts and hypertrophic chondrocytes during bone formation, but the mutant Runx2 exhibited reduced expression in hypertrophic chondrocytes and partially impaired nuclear localization, resulting in arrested osteoblast and chondrocyte maturation. Our mutant mouse model provides a valuable *in vivo* platform to study CCD pathogenesis, mechanisms of tooth root furcation, and therapeutic interventions targeting dysfunctional RHD.

## Introduction

Cleidocranial dysplasia (CCD; OMIM #119600) is a rare autosomal dominant skeletal and dental disorder caused by haploinsufficiency of the *Runt-related transcription factor 2* (*RUNX2*)^(1,2)^. CCD affects one in a million people worldwide, with no gender or ethnic differences, although mild cases may be underdiagnosed^(3)^. Clinical features include clavicular hypoplasia or aplasia, midfacial hypoplasia, delayed closure of cranial sutures, Wormian bones, and a persistent anterior fontanelle closure, often with short stature and dental anomalies^(1)^. Prolonged retention of deciduous teeth and multiple unerupted supernumerary permanent teeth are frequently encountered in dental clinics^(3,4)^, requiring long-term multidisciplinary care, including oral and maxillofacial surgery and orthodontics, to manage function, aesthetics, and patient burden.

The human *RUNX2* and mouse *Runx2* genes comprise 8 exons, spanning approximately 218 kb (hg38, NM_001024630.4) and 206 kb. (mm39, NM001271627.2), respectively^(5,6)^. Homozygous *Runx2*-deficient mice die shortly after birth from respiratory failure, exhibiting a complete lack of ossification and arrested embryonic tooth development, while heterozygous mice exhibit persistent open anterior fontanelle, clavicle hypoplasia, and other skeletal abnormalities resembling human CCD^(2,7)^. RUNX2/Runx2 is a key transcription factor in skeletal development, regulating mesenchymal commitment to the osteoblastic lineage, proliferation and maturation of chondrocytes and osteoblasts, and transdifferentiation of hypertrophic chondrocytes into osteoblasts^(8)^. Intramembranous ossification strongly depends on *Runx2* and requires higher levels of its expression compared to endochondral ossification^(5)^. In *Runx2^+/-^*mice, as in human CCD, delayed eruption of incisors and maxillary first molars occurs due to impaired osteoclast recruitment in the alveolar bone, although neither impacted nor supernumerary teeth are present^(9)^.

RUNX2 contains several functional domains, including a glutamine alanine-rich domain (QA), a highly conserved Runt homology domain (RHD), which is also referred to as the Runt domain, a proline serine threonine domain (PST), and a C-terminal VWRPY motif, and harbors a nuclear transition signal (NLS) at the C-terminus of the RHD and a nuclear matrix targeting signal (NMTS)^(10)^. More than 200 *RUNX2* mutations have been reported in CCD patients, including frameshift, missense, nonsense, and splice-site variants^(6,11)^. While nonsense and frameshift mutations are distributed throughout the gene, most missense mutations cluster within the 128-amino acid RHD, which is critical for DNA binding and heterodimerization with core-binding factor subunit β (Cbfb)^(6,12,13)^. Many pathogenic mutations associated with CCD, especially those linked to severe dental anomalies, are missense or nonsense mutations affecting the RHD^(6)^. However, the mechanisms by which these mutations cause severe CCD have long remained unclear, owing to the absence of *in vivo* models with RHD-specific mutations and the limitations of cellular systems that cannot fully recapitulate the disease complexity.

Recently, several disease-specific induced pluripotent stem (iPS) cell lines carrying mutations such as p.R225Q, p.S124W, p.R391X, and p.Q67X have been established from patients^(14–16)^. iPS cells harboring nonsense mutations p.Q67X in the QA domain and p.R391X in the NTMS domain exhibit reduced osteogenic differentiation capacity^(14)^. Transplantation of osteoblasts induced from p.Q67X iPS cells into rat calvarial defects led to limited bone regeneration, which was rescued by genetic correction of the *RUNX2* mutation^(14)^. While disease-specific iPS cells are useful for modeling altered differentiation potential and cellular behavior *in vitro* but fail to recapitulate morphogenetic processes during development. Furthermore, iPS cells from CCD patients with heterozygous *RUNX2* mutations do not allow comprehensive evaluation of mutant protein function. Despite extensive studies on RUNX2 and CCD, the *in vivo* impact of RHD mutations is still not well characterized.

In this study, we generated two novel mouse models carrying RHD mutations: a missense mutation (c.695G>A; p.R232Q) and a frameshift mutation (c.697_698delGA; p.E233TfsTer9). These models serve as a robust platform for investigating CCD pathogenesis and uncover previously unrecognized abnormalities in tooth root furcation. By enabling detailed functional analysis of RHD mutations in a disease-relevant context, they provide new insights into CCD pathogenesis and may contribute to the development of targeted therapies.

## Materials and methods

### Animals

C57BL/6 mice were purchased from CLEA Japan, Inc. (Tokyo, Japan) and bred for experiments. Homozygous *Runx2^m/m^* and *Runx2^112/112^* mice exhibited perinatal lethality, which was expected^(7,12)^, and no intervention was undertaken as it was unpreventable. Mice were maintained under a 12-hour light/dark cycle (lights on from 8:00 to 20:00) at 23 ± 3 °C and 50

± 10% humidity. Animals were provided γ-irradiated MF chow (Oriental Yeast Co., Ltd., Tokyo, Japan) and autoclaved water ad libitum. All procedures were approved by the Animal Care Committee of the Institute for Frontier Life and Medical Sciences, Kyoto University, as well as the Committee of Animal Experimentation and the Recombinant DNA Committee of Hiroshima University.

### Generation of TALEN-mediated *Runx2* mutant mouse lines

TALEN plasmids targeting mouse *Runx2* were constructed using the Platinum Gate TALEN Kit (Addgene, Watertown, MA) as previously described^(17)^. After linearization of TALEN plasmids for *Runx2-TALEN-R* and *-L* with SmaI, mRNAs were synthesized using the mMESSAGE mMACHINE T7 ULTRA Kit (Thermo Fisher Scientific, Waltham, MA) and purified with the MEGAclear kit (Thermo Fisher Scientific). The TALEN mRNAs and an ssODN carrying the p.R232Q and a silent ApaI site were microinjected into C57BL/6 oocytes. Injected eggs were transferred into pseudopregnant ICR females. Genotyping was performed by PCR on genomic DNA extracted from earpieces using specific primers (Table S3). The products were digested with SmaI and ApaI, and confirmed by direct sequencing using the BigDye Terminator Cycle Sequencing kit and an ABI 3100 Genetic Analyzer (Applied Biosystems, Thermo Fisher Scientific). Fragments were analyzed by agarose gel electrophoresis or a microchip electrophoresis system for DNA/RNA analysis (MultiNA, SHIMADZU, Kyoto, Japan).

### Skeletal preparation

Whole-mount skeletal staining was performed as previously described^(18)^. Embryos or mice were skinned, fixed in ethanol, stained with Alcian blue 8GX (Sigma-Aldrich, Merck KGaA, Darmstadt, Germany), washed with ethanol, and cleared in 1% KOH. Samples were then stained with 0.05% Alizarin Red S (Sigma-Aldrich) in 1% KOH, followed by additional clearing in 1% KOH. Images were acquired using a MZ16 FA stereomicroscope equipped with a DFC7000 T camera (Leica Microsystems (Leica), Wetzlar, Germany).

### Tissue sections

For frozen sections, embryos were fixed in 4% paraformaldehyde (PFA) in phosphate buffered saline (PFA/PBS), infiltrated with 20% sucrose/PBS, embedded in Tissue Tek O.C.T compound (Sakura Finetek, Tokyo, Japan), sectioned at 4-6 µm. For paraffin sections, embryos were fixed in 4% PFA/PBS, decalcified with Morse’s solution^(19)^, dehydrated, immersed in methyl benzoate (Nacalai Tesque, Kyoto, Japan), embedded in paraffin (Thermo Fisher Scientific), sectioned at 6 µm.

### Histological staining

Frozen sections were stained with ALP substrate solution of TRAP/ALP Stain Kit (FUJIFILM Wako Pure Chemical Corporation (FUJIFILM Wako), Osaka, Japan), or stained with 1% alizarin red (Muto Pure Chemicals, Tokyo, Japan). For toluidine blue staining, deparaffinized and hydrated sections were stained with 0.05% toluidine blue solution (pH 4.1; Muto Pure Chemicals).

### *In situ* hybridization

Digoxygenin (DIG)- labelled RNA probes for *Col2a1*^(20)^, *Acan*^(20)^, *Ihh*, *Col10a1*^(21)^, *Runx2*^(22)^, *Spp1*^(23)^ and *Col1a1*^(24)^ were synthesized using a DIG RNA labelling kit (F. Hoffmann-La Roche Ltd. (Roche), Basel, Switzerland). Mouse *Col2a1* and *Ihh* cDNA were amplified by RT-PCR using the following primers (Table S3). Paraffin sections were deparaffinized, rehydrated and treated with 10 µg/mL Proteinase K (Roche), followed by post-fixation with 4% PFA/PBS for 10 min. Sections were hybridized with DIG-labeled RNA probes at 65°C for 16 h, washed, and incubated with anti-DIG Fab fragment conjugated with alkaline phosphatase (anti-DIG-AP; Roche) and BM purple (Roche). Counterstaining was performed with eosin (Sakura Finetek).

### μCT imaging

Mice were anesthetized with a combination of medetomidine, midazolam, and butorphanol (0.1 mL per 10 g body weight). Calvaria and clavicle were scanned with SkyScan1176 (Bruker, Antwerp, Belgium; 17.5 µm resolution, 50 kV and 500 µA), and teeth were scanned with CosmoScan GXIII (Rigaku Holdings Corporation, Tokyo, Japan; 12.9 µm resolution, 100 kV and 120 µA). 3D rendering was performed using Amira software (Thermo Fisher Scientific, version 2021.1).

### Plasmid construction

Mouse Runx2 cDNA was amplified from *pSG5-mRunx2* and cloned into the EcoRI site of *pcDNA-FLAG*^(20)^ to generate *pcDNA3 Runx2 WT* using primers. Mutant vectors carrying the p.R232Q, p.R232W, or p.R200X mutations (*pcDNA3 Runx2 RQ*, *RW*, and *RX*, respectively) were generated by inverse-PCR with KOD Plus DNA polymerase (TOYOBO, Osaka, Japan) using *pcDNA3 Runx2 WT* as the template. See Table S2 for primers.

### Dual luciferase reporter assay

HEK293T cells (Takara Bio Inc., Shiga, Japan) were cultured in Dulbecco’s modified Eagle’s medium with high glucose (Sigma-Aldrich) supplemented with 10% heat-inactivated-FBS (Hana-Nesco Bio Corp., Osaka, Japan), 2 mM L-glutamine (FUJIFILM Wako), and 50 mg/mL kanamycin (Sigma-Aldrich) at 37°C in 5% CO_2_. Cells were transiently transfected with *p6OSE2-luc*^(25)^ and *pGL4.10[luc2]*, *pGL4.74[hRluc/TK]* (Promega, Madison, WI), and *pcDNA3 Runx2 WT or mutant* vectors. Transfection was performed using Lipofectamine LTX Reagent or Lipofectamine 3000 Reagent (Thermo Fisher Scientific). After 48 h, luciferase activities were measured using the PicaGene Dual Sea Pansy Luminescence Kit (Toyo ink, Tokyo, Japan) and Centro XS^3^ LB 960 (Berthold Technologies, Bad Wildbad, Germany).

### Western blotting

Forelimbs were frozen in liquid nitrogen, crashed using SK mill (SK-200, Tokken,Inc, Chiba, Japan), and lysed in 8 M urea extraction buffer [8 M urea, 50 mM Tris-HCl (pH 8.0), 1 mM DTT and 1 mM EDTA]. HEK293T cells were transfected with WT or mutant Runx2 vector using Lipofectamine 3000 Reagent (Thermo Fisher Scientific), and lysed in 8 M urea extraction buffer. Protein concentrations were determined using BCA Protein Assay Kit (Thermo Fisher Scientific), and samples were mixed with 6x SDS sample buffer (Nacalai tesque, Inc., Kyoto, Japan), boiled, and loaded onto a 10% SDS-polyacrylamide gels (e-PAGEL, ATTO, Tokyo, Japan). Samples were separated by SDS-PAGE and transferred to PVDF membranes using iBlot3 (Invitrogen). Membrane were blocked with 3% skim milk (FUJIFILM Wako) in Tris-buffered saline with 0.05% Tween 20 (TBS-T), incubated with rabbit anti-Runx2 monoclonal antibody (1:1250, Cell Signalling Technology, Danvers, MA) and mouse anti-β-actin monoclonal antibody (1:1250, FUJIFILM Wako) overnight, then with IRDye 680RD Goat anti-Rabbit IgG and IRDye 800CW Goat anti-Mouse IgG (1:15000, LI-COR, Lincoln, NE). Bands were visualized using Odyssey CLx (LI-COR).

### Immunostaining

Frozen sections were subjected to antigen retrieval in sodium citrate buffer (pH6.0) by microwave heating, followed by permeabilization with 0.2% TritonX-100 (Sigma-Aldrich). After blocking with 2% skim milk/PBS, sections were incubated with rabbit anti-Runx2 monoclonal antibody (1:1600, Cell Signaling Technology) or rabbit anti-Collagen X monoclonal antibody (1:500, Abcam, Cambridge, UK) overnight, washed with PBS, and incubated with Goat anti-Rabbit IgG (H+L) Highly Cross-Adsorbed Secondary antibody conjugated with Alexa Fluor Plus 594 (1:500, Thermo Fisher Scientific). Nuclei were counterstained with 4’,6-diamidino-2-phenylindole (DAPI) (Sigma-Aldrich). Stained sections were mounted using VECTASHIELD Hard-Set Mounting Medium (Vector Laboratories, Inc., Newark, CA). Images were acquired using DMRXA microscope with DFC310 FX Camera and Stellaris 5 confocal laser scanning microscope (Leica).

### Image analysis

To analyze Runx2 nuclear localization, maximum projection images were generated from confocal images using LAS X software (Leica). Cells in the Col10^+^ hypertrophic cartilage layer were analyzed with ImageJ software (National Institutes of Health, Bethesda, MD). Regions of interest (ROIs) for whole-cell Runx2 [ROIs(r)] were defined from Runx2 signals using thresholding based on the Huang algorithm^(26)^. Nuclear regions of interest [ROIs(d)] were defined from DAPI images with thresholds set using the Default (IsoData) algorithm^(27)^, excluding ROIs smaller than 50 pixels. ROIs(r) and ROIs(d) were overlayed on Runx2 images, and the respective Runx2 intensities in whole cells and nuclei were measured using Integrated Density. Nuclear localization was calculated as: %Nuclear = (Runx2 intensity in nucleus/Runx2 intensity in the whole cell) ×100. Only cells with overlapping ROI(r) and ROI(d), confirmed by visual inspection, were included. Multiple ROI(d) within one ROI(r) were summed.

### Statistical analysis

Statistical significance between two groups was determined using Student’s *t*-test or Mann-Whitney U test. Comparisons among multiple groups were performed by one-way analysis of variance (ANOVA) followed by Tukey’s multiple comparison test. Graphpad Prism 10 (Dotmatics, Boston, MA, version 10. 2. 3) was used.

## Results

### Establishment of CCD mouse models harboring *Runx2* missense or frameshift mutations

The mouse *Runx2* gene, located on chromosome 17 and composed of 8 exons, is transcribed from two distinct promoters: distal P1 and proximal P2, generating isoforms with different N-terminal amino acid sequences^(6)^ (Fig. 1A, Fig. S1A). The 128-amino-acid RHD crucial for DNA binding, is encoded by exons 2 to 4, and the NLS by exons 4 and 5 (Fig. 1B). CCD patients with a heterozygous p.R225Q mutation in human *RUNX2* show multiple supernumerary teeth and membranous bone dysplasia^(15,28–30)^. In mouse *Runx2*, this mutation corresponds to p.R232Q owing to seven additional amino acids at the N-terminus. We introduced the c.695G>A (p.R232Q) mutation into the RHD of *Runx2*. To introduce the mutation via homology-directed repair (HDR), we used Platinum TALENs and a single-stranded oligodeoxynucleotide (ssODN) containing the c.695G>A substitution and a silent ApaI site for genotyping (Fig. 1a). Microinjection of Platinum TALEN mRNAs and a designed ssODN into fertilized eggs generated 51 mice, including 14 with non-homologous end joining induced indels and 2 carried the HDR-mediated knock-in allele (Table S1). Lines carrying the missense (*m*) or the double-nucleotide deletion (*Δ2*) mutation were established from founder mice (Fig. 1C, Fig. S1A, B). In *Δ2* mutants, deletion of nucleotides 697G and 698A caused a frameshift mutation, generating a premature stop codon at 241 (Fig. S1B, Fig. 1C). For genotyping, SmaI digestion yielded 115 bp and 80 bp fragments for the wild-type allele, while mutant alleles produced an undigested 195 bp fragment, and ApaI digestion generated 113 bp and 82 bp fragments specific to the missense mutant allele (Fig. S2). Direct sequencing confirmed that *Runx2^m/+^*carried both the c.690T>G silent andc.695G>A missense mutations, while *Runx2^112/+^* harbored a two-nucleotide deletion causing a premature stop codon. (Fig. 1D). Thus, we successfully established two distinct CCD model mice: one with a clinically relevant RHD missense mutation associated with severe CCD and another with a frameshift mutation.

**Fig. 1.**
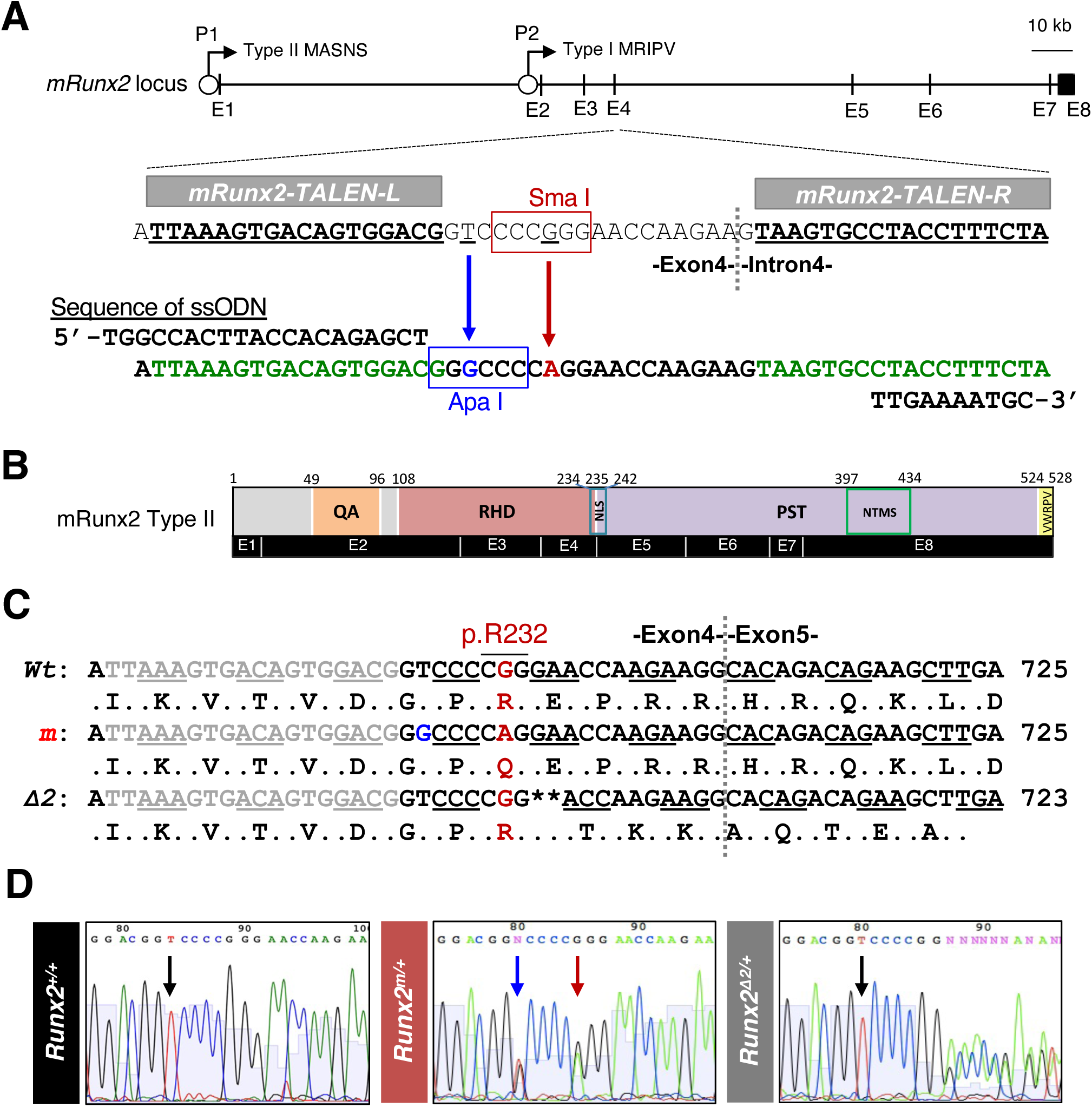
Generation of *Runx2* mutant mice using Platinum TALENs. (A) Schematic representation of the mouse *Runx2* gene with eight exons (labeled E1-E8 below), distal (P1) and proximal (P2) promoters for Type II (MASNSL) and Type I (MRIVD) isoforms, and target sites of Platinum TALENs (*mRunx2-TALEN-L* and *-R*). A 10 kb scale bar is shown. The SmaI site (red box) includes G695 (underlined). The ssODN contains target sites (green), a T>G silent mutation at 690 creating an ApaI site (blue box), and a G>A mutation at 695 disrupting the SmaI site. (B) Domain structure of the mRunx2 Type II protein (528 aa) and its corresponding exons. The protein comprises a QA repeat domain, RHD, NLS, PST-rich domain, NMTS, and a C-terminal VWRPY motif. Amino acid positions are indicated above, and the corresponding exons (E1-E8) are shown below. (C) Nucleotide sequences are shown for the wild-type (*Wt*, 670-725), missense mutant (*m*, c.695G>A; p.R232Q, 670-725), and two-base deletion (*β2*, c.697_698delGA; p.E233TfsTer9, 670-723) alleles. *Wt* is on top, *m* in the middle, and *β2* at the bottom. (D) Representative electropherograms of the targeted *Runx2* region in *Runx2^+/+^*, *Runx2^m/+^*, and *Runx2^112/+^* mice. A black arrow marks T at 690 in *Runx2^+/+^*and *Runx2^m/+^* mice. A blue arrow indicates overlapping T and A peaks at 690 in *Runx2^m/+^* mice, and a red arrow shows overlapping A and G peaks at position 695 in *Runx2^m/+^* mice.

### Defective bone formation in homologous *Runx2* RHD-mutant mice

Homozygous *Runx2^m/m^* and *Runx2^112/112^* mice exhibited perinatal lethality, likely due to respiratory failure caused by severe skeletal abnormalities, which was predictable based on previous studies of *Runx2* knockout mice^(7,12)^. At embryonic day 18.5 (E18.5), *Runx2^+/+^*, *Runx2^m/+^*, and *Runx2^112/+^* embryos were similar in size and morphology, while *Runx2^m/m^* and *Runx2^112/112^* embryos were smaller with limb shortening, facial underdevelopment, and calvarial defects (Fig. S3A-E). Skeletal development at E18.5 was compared among wild-type, heterozygous, and homozygous *Runx2* RHD mutants (Fig. 2A-J). In *Runx2^+/+^*embryos, intramembranous ossification progressed in the calvaria and clavicles, and endochondral ossification in the axial and appendicular skeletons (Fig. 2A, F, G). In *Runx2^m/+^* or *Runx2^112/+^* embryos, intramembranous ossification occurred but was impaired, causing clavicle hypoplasia and delayed fontanelle closure (Fig. 2B, C, H, I). In *Runx2^m/m^* or *Runx2^112/112^* embryos, intramembranous ossification was absent, causing skull and clavicle defects, and endochondral ossification was impaired, with long bones remaining mostly cartilaginous and mineralization limited to the forearms and the lower legs (Fig. 2D, E, J). Forelimb calcification was more delayed in *Runx2^m/m^*than in *Runx2^112/112^* embryos (Fig. 2D, E).

**Fig. 2.**
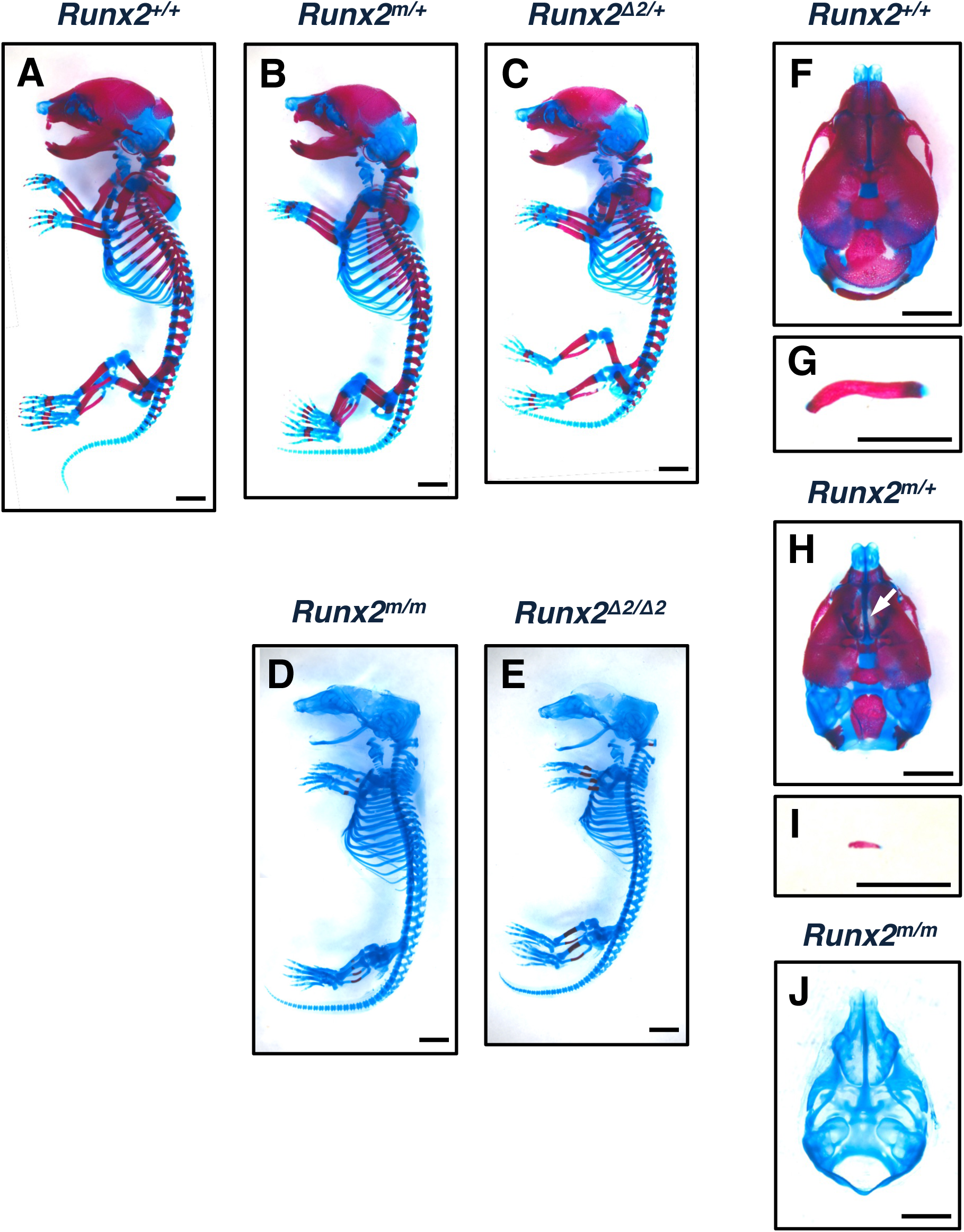
Skeletal phenotypes of *Runx2* mutant mice. (A-J) Skeletal preparations of *Runx2^+/+^* (A, F, G), *Runx2^m/+^* (B, H, I), *Runx2^112/+^* (C), *Runx2^m/m^*(D, J), and *Runx2^112/112^* (E) mice at E18.5. Lateral views of embryos (A-E), clavicles (G, I) and dorsal views of the calvaria (F, H, J) are presented. A white arrow indicates weak calcification. Scale bars, 2 mm.

Histological sections of the forearm from *Runx2^+/+^*, *Runx2^m/+^*, *Runx2^m/m^* and *Runx2^112/112^* embryos at E18.5 were stained with toluidine blue (TB), alkaline phosphatase (ALP), and Alizarin Red (AR) (Fig. 3A-C). In *Runx2^+/+^* and *Runx2^m/+^* embryos, primary ossification canters formed in the diaphysis, with metachromatic cartilage present in the metaphysis and epiphysis. In *Runx2^m/m^* and *Runx2^112/112^* embryos, the entire bone primordium consisted exclusively of cartilage (Fig. 3A). ALP activity was high from the diaphysis to the metaphysis in *Runx2^+/+^* and *Runx2^m/+^* embryos, but was restricted to the hypertrophic/calcified cartilage in the center of the ulna and radius of *Runx2^m/m^* and *Runx2^112/112^* embryos (Fig. 3B). Although the calcified area was smaller than the ALP^+^ area in all genotypes, in *Runx2^m/m^* and *Runx2^112/112^* embryos, both regions were markedly smaller than those in *Runx2^+/+^* and *Runx2^m/+^*embryos and were confined to the central diaphysis where hypertrophic chondrocytes were present (Fig. 3C). Notably, calcified cartilage was smaller in *Runx2^m/m^* than in *Runx2^Δ2/Δ2^* embryos, suggesting delayed cartilage maturation (Fig. 3A-C).

**Fig. 3.**
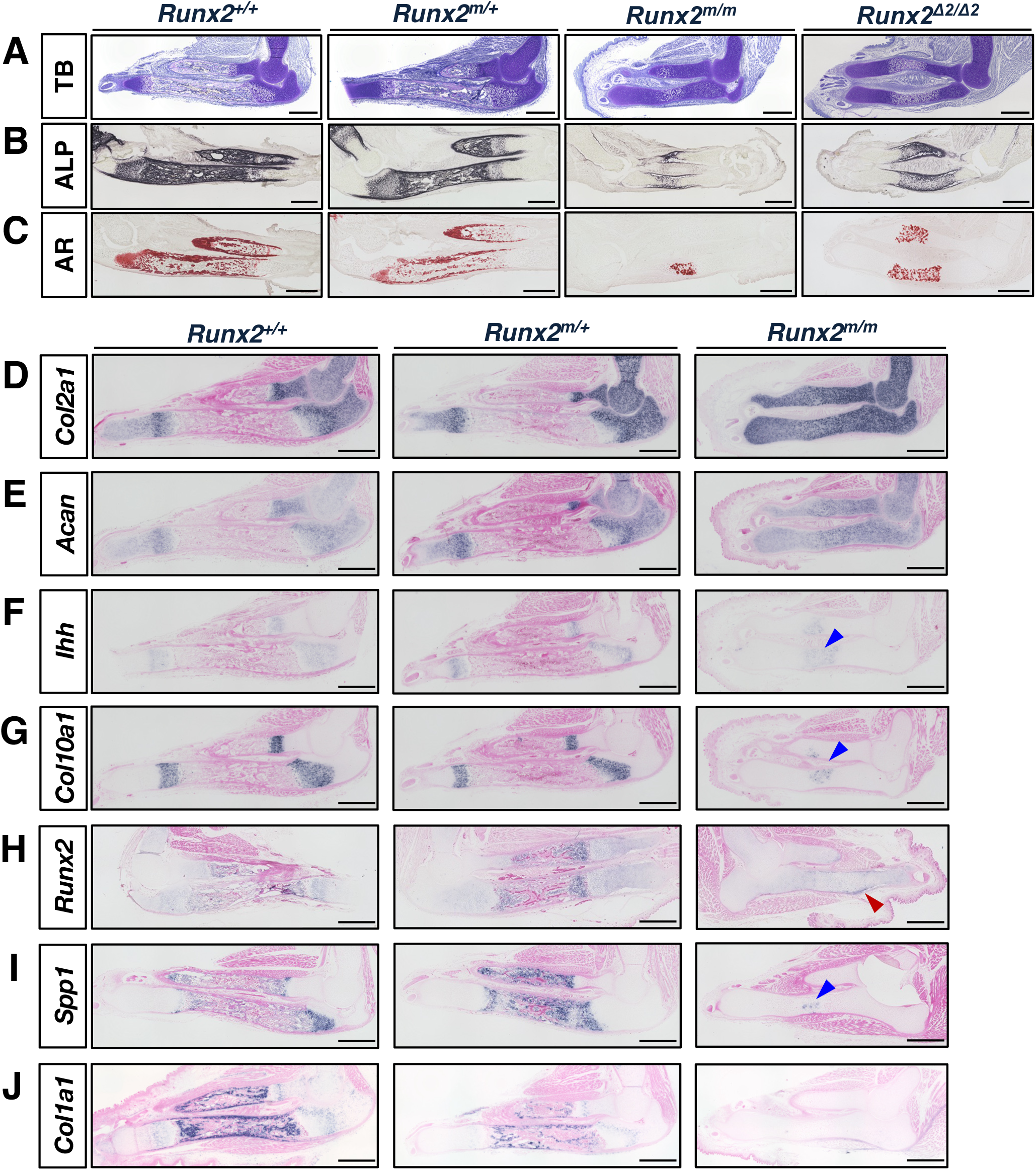
Histology and *in situ* hybridization of forearms in wild-type and *Runx2* mutant mice. (A-C) TB, ALP, and AR staining of forearms prepared from *Runx2^+/+^*, *Runx2^m/+^*, *Runx2^m/m^* and *Runx2^112/112^* embryos at E18.5. (D-J) Expression of *Col2a1* (D), *Acan* (E), *Ihh* (F), *Col10a1* (G), *Runx2* (H), *Spp1* (I), and *Col1a1* (J) in the forearm of *Runx2^+/+^*, *Runx2^m/+^*, or *Runx2^m/m^* embryos at E18.5. Blue arrowheads in (F), (G), and (I) mark prehypertrophic/hypertrophic chondrocytes. A red arrowhead in (H) indicates the perichondrium. Scale bars, 500 µm.

To assess chondrogenic and osteogenic marker expression, the forearm of *Runx2^+/+^*, *Runx2^m/+^* and *Runx2^m/m^*embryos at E18.5 were analyzed by *in situ* hybridization (Fig. 3D-J). Early chondrogenic markers *Col2a1* and *Acan* were expressed in proliferating and mature chondrocytes in *Runx2^+/+^*, *Runx2^m/+^*, and *Runx2^m/m^* embryos (Fig. 3D, E). *Ihh* was expressed in prehypertrophic chondrocytes in *Runx2^+/+^* and *Runx2^m/+^*embryos, whereas its expression was only faintly detected in the central diaphysis of *Runx2^m/m^* embryos (Fig. 3F). *Col10a1* was expressed in hypertrophic chondrocytes of the metaphysis in *Runx2^+/+^* and *Runx2^m/+^* embryos, but only weakly detected in hypertrophic chondrocytes in the central diaphysis in *Runx2^m/m^* embryos (Fig. 3G). In *Runx2^+/+^* and *Runx2^m/+^* embryos, *Runx2* and *Spp1* were expressed in osteoblasts, osteocytes, and chondrocytes (Fig. 3H, I). In *Runx2^m/m^* embryos, *Runx2* was faintly expressed in chondrocytes and perichondrial cells, while *Spp1* was detected in a small population of hypertrophic chondrocytes in the diaphysis (Fig. 3H, I). *Col1a1* was detected in osteoblasts, osteocytes, cells of the periosteum and perichondrium in *Runx2^+/+^*and *Runx2^m/+^* embryos, but only faintly in the perichondrium of *Runx2^m/m^* embryos (Fig. 3J).

These findings demonstrate that RHD mutations disrupt both intramembranous and endochondral ossification, emphasizing the essential role of the RHD in skeletal development.

Functional impairment of Runx2 caused by the p.R232Q mutation in the RHD

Wild-type and p.R232Q mutant Runx2 proteins were analyzed by Western blotting using whole-cell extracts from HEK293T cells transfected with each construct (Fig. 4A). Immunoblotting with an anti-RUNX2 antibody recognizing a peptide around Arg (R) at 267 of human RUNX2 confirmed the expression of both wild-type and mutant Runx2, with no apparent differences, although a sight size shift was observed due to the FLAG tag (Fig. 4A, Fig. S4A). Next, Western blotting with the same anti-RUNX2 antibody was performed on forelimb extracts from *Runx2^+/+^*, *Runx2^m/+^*, and *Runx2^m/m^* embryos at E18.5 to assess Runx2 expression (Fig. 4B, Fig. S4B). The reported molecular weights are approximately 57 kDa for isoform 1 (MASNS) and 56 kDa for isoform 2 (MRIPV)^(31)^. Both wild-type and p.R232Q mutant Runx2 showed similar sizes in HEK293T whole-cell extracts transfected with each construct (Fig. 4A, Fig. S4A) and in tissue extracts from *Runx2^+/+^*, *Runx2^m/+^*and *Runx2^m/m^* embryos, although the expression level was markedly reduced in *Runx2^m/m^* embryos compared to *Runx2^+/+^* embryos (Fig 4B, Fig. S4Bb).

**Fig. 4.**
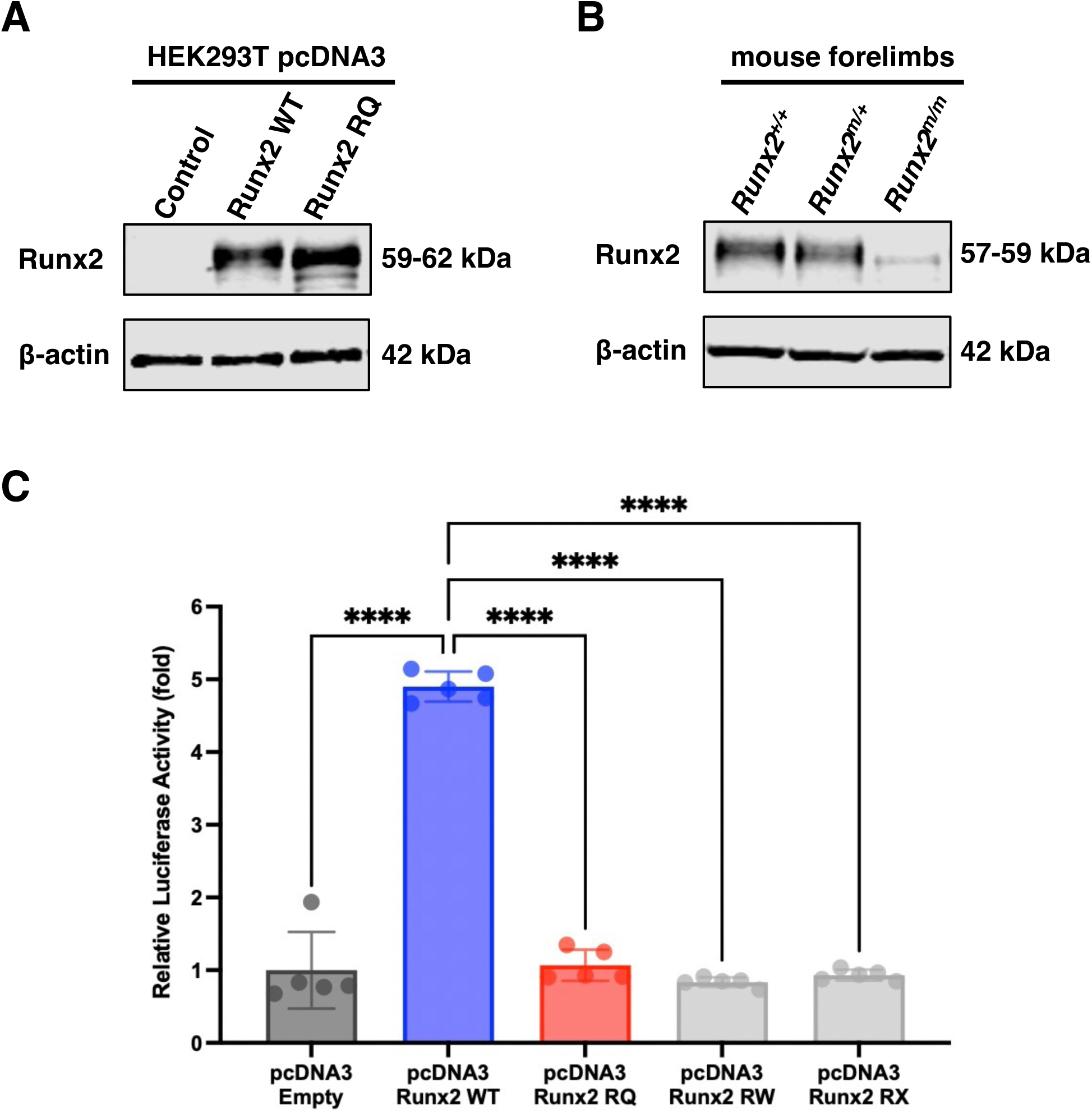
Expression and transcriptional activity of wild-type and mutant Runx2. (A) Expression of wild-type and missense mutant Runx2 in HEK293T cells. Cells were transfected with an empty vector as the control, wild-type *Runx2,* or the *Runx2 RQ* mutant construct, all using the pcDNA3 vector. (B) Expression of Runx2 in forelimb extracts from *Runx2^+/+^*, *Runx2^m/+^* and *Runx2^m/m^* mice at E18.5. β-Actin was used as a loading control in both (A) and (B). (C) Dual luciferase assay in HEK293T cells co-transfected with the *p6OSE2-luc* reporter vector and either empty vector, *Runx2 WT, Runx2 RQ, RW, or RX* mutant constructs using the *pcDNA3 vector*. *pGL4.74[hRluc/TK]* was also co-transfected for normalization. Values for each of five wells were normalized to *pGL4.74[hRluc/TK]* and are shown as fold induction relative to the empty vector. Data present one of ≧3 independent experiments (mean ± SD, n = 3). \*\*\*\**P < 0.0001 vs. Runx2 WT*.

The *p6OSE2-luc* vector contains six tandem repeats of osteoblast-specific element 2 (OSE2, AACCACA) from the mouse osteocalcin promoter, a Runx2 binding site, followed by a minimal promoter driving firefly luciferase expression, allowing quantification of Runx2-dependent transcription^(25)^. To evaluate the effect of RHD mutations, HEK293T cells were transfected with wild-type or mutant Runx2 (p.R232Q, p.R232W, or p.R200X) (Table S2), along with *p6OSE2-luc* and the *pGL4.74[hRlux/TK]* internal control, and dual luciferase assays were performed (Fig. 4C). Wild-type Runx2 showed a 6.7-fold increase in transcriptional activity compared to the empty vector, whereas the p.R232Q, p.R232W, or p.R200X mutants exhibited activity comparable to the empty vector, indicating that missense and nonsense mutations in the RHD abolish Runx2-mediated activation via OSE2 (Fig. 4C).

The p.R232Q mutation lies near the NLS in the C-terminal region of the RHD^(32)^, potentially affecting Runx2 nuclear translocation. To examine the impact of the mutation on intracellular localizations, immunostaining was performed on sections of the ulna sections from *Runx2^+/+^*and *Runx2^m/m^* embryo at E18.5 using anti-Col10 and anti-Runx2 antibodies (Fig. 5A-D). Consistent with the results of *in situ* hybridization (Fig. 3G, H), In *Runx2^+/+^* embryos, Col10 accumulated abundantly in the metaphysis and in cartilage remnants of the primary spongiosa (Fig. 5A), while Runx2^+^ cells were distributed from the metaphysis to the diaphysis (Fig. 5B). In *Runx2^m/m^* mice, Col10^+^ hypertrophic chondrocytes were confined to the central diaphysis (Fig. 5C), and Runx2^+^ cells were more widely distributed but with markedly weaker signal intensity (Fig. 5D). In hypertrophic chondrocytes of *Runx2^+/+^* embryos, Runx2 was predominantly localized in the nucleus (Fig. 5E, F). In contrast, many cells in *Runx2^m/m^* embryos exhibited partial nuclear translocation, but cytoplasmic localization was also observed, and overall nuclear localization of Runx2 was significantly reduced (Fig. 5G, H). Quantification of the nuclear-to-total Runx2 fluorescence intensity ratio revealed a significant decrease in *Runx2^m/m^* embryos compared to *Runx2^+/+^* embryos (Fig. 5I, Fig. S5).

**Fig. 5.**
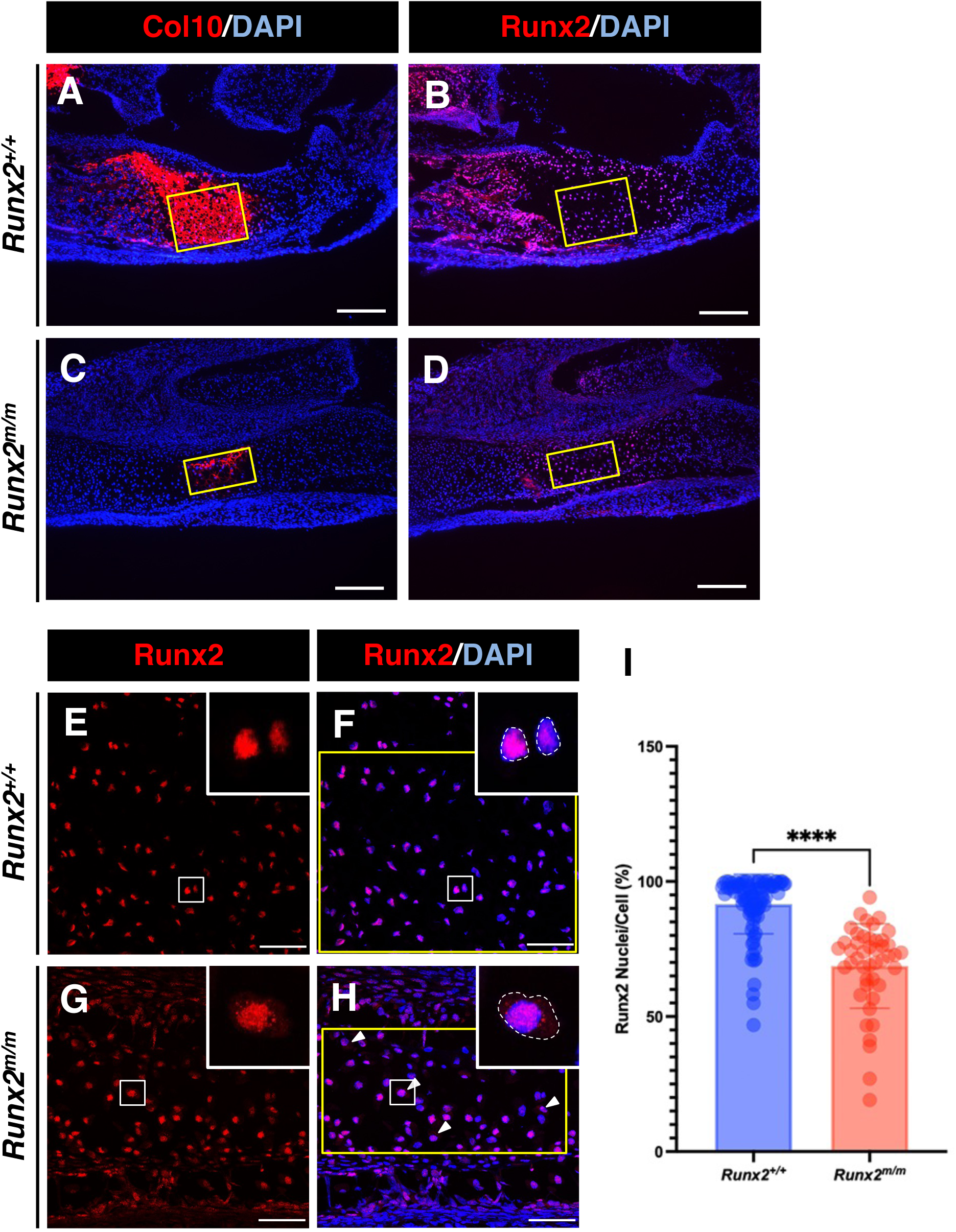
Altered expression and subcellular localization of mutant Runx2 in chondrocytes. (A-D) Immunostaining of Col10 (A, C) and Runx2 (B, D) proteins in the ulna of *Runx2^+/+^* and *Runx2^m/m^* embryos at E18.5. Nuclei are counterstained with DAPI. Yellow boxes in (A) and (**C**) indicate the Col10^+^ prehypertrophic/hypertrophic cartilage zone. The corresponding regions to them are shown in the semi-serial sections in (B) and (D). The number of cells in the yellow boxes of (A) and (C) indicating Col10^+^ region differs between *Runx2^+/+^* and *Runx2^m/m^*. (E-H) Confocal images of wild-type and mutant Runx2 (red) and nuclei (blue) in prehypertrophic/hypertrophic chondrocytes. Representative localization patterns of wild-type (E, F) and mutant (G, H) Runx2 are highlighted by rectangles and shown in insets. (I) Nuclear localization rate of Runx2 in chondrocytes of the ulna in *Runx2^+/+^*and *Runx2^m/m^* embryos. Runx2 signal intensities in the whole-cell (dashed regions in f and h) and those in nucleus were quantified using imageJ. Data are means ± SD (*Runx2^+/+^*, n = 88 cells; *Runx2^m/m^*, n = 45 cells). *\*\*\*\*P < 0.0001* (Mann-Whitney *U*-test). Scale bars, 200 µm (A-D), 50 µm (E-H).

These results indicate that the p.R232Q mutation in the RHD impairs Runx2 function in skeletal development by reducing its expression, transcriptional activation capacity, and nuclear localization.

### Postnatal bone abnormalities recapitulating human CCD in *Runx2^m/+^*mice

*Runx2^m/+^* mice showed smaller body size and length at 1 month of (Fig. 6A) and 18.3% lower body weight at 4 months compared to *Runx2^+/+^*mice (Fig. 6B). Three-dimensional (3D) rendering images from micro-CT (µCT) scans revealed significantly smaller clavicles in *Runx2^m/+^* mice (Fig. 6C), and skeletal preparations at 3 months confirmed markedly hypoplastic clavicles and scapulae compared to *Runx2^+/+^* mice (Fig. 6D).

**Fig. 6.**
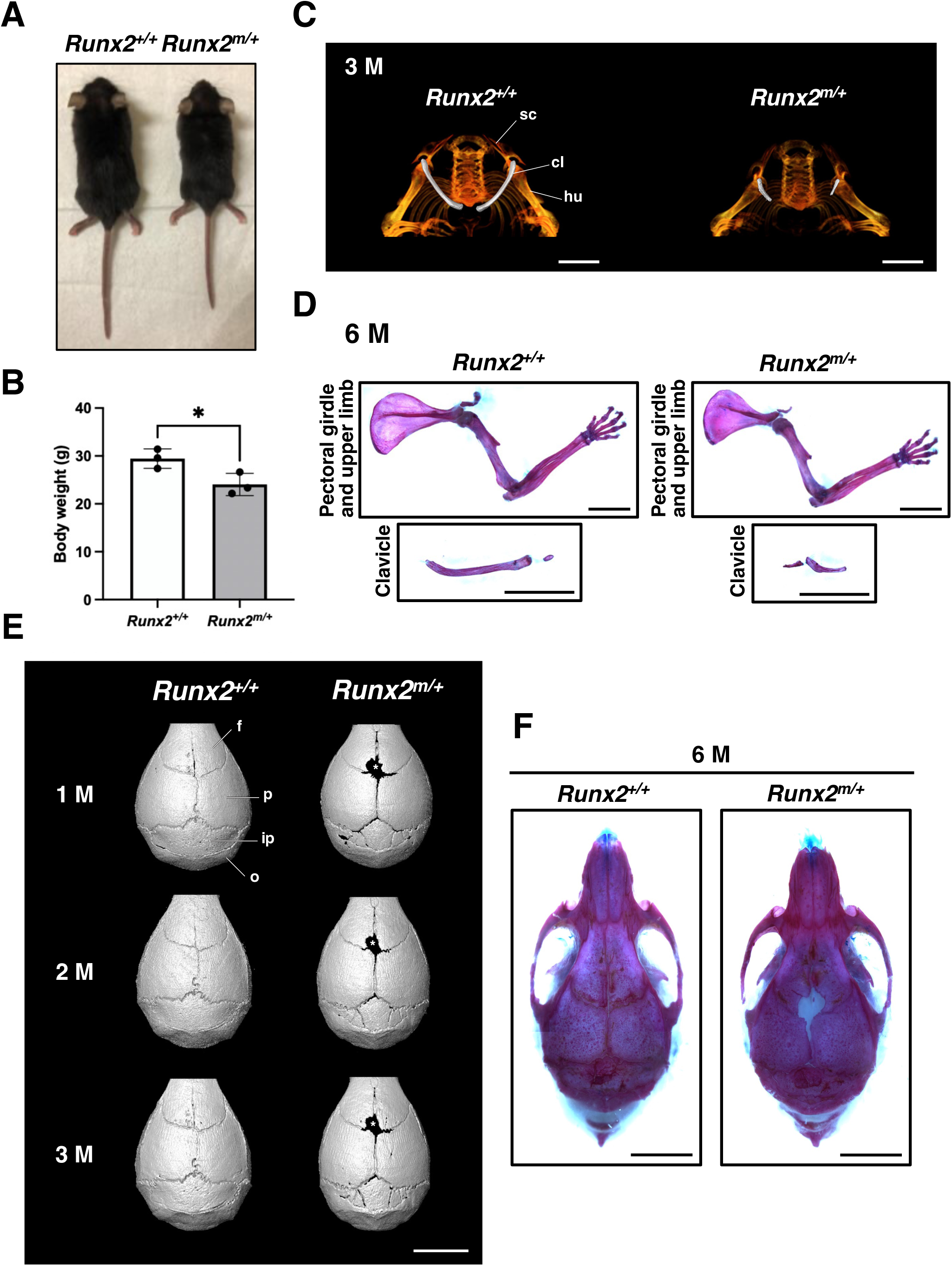
Postnatal skeletal development in *Runx2^+/+^* and *Runx2^m/+^* mice. (A) Gross appearance of *Runx2^+/+^* and *Runx2^m/+^* mice at P30. (B) Body weight of *Runx2^+/+^* and *Runx2^m/+^* mice at 16 weeks (n = 3). *\*P < 0.05 vs. Runx2^+/+^* (unpaired Student’s *t*-test). (C) µCT-based 3D reconstruction images of the axial and appendicular skeleton in the upper body of 3-month-old *Runx2^+/+^* and *Runx2^m/+^* mice. (D) Skeletal preparations of the pectoral girdle and upper limb in *Runx2^+/+^* and *Runx2^m/+^* mice at 6 months. € µCT-based 3D reconstruction images of the calvaria in *Runx2^+/+^*and *Runx2^m/+^* mice at 1, 2, and 3 months. (F) Skeletal preparations of the skull from *Runx2^+/+^* and *Runx2^m/+^*mice at 6 months. Abbreviations: sc, scapula; cl, clavicle; hu, humerus; f, frontal bone; p, parietal bone; ip, interparietal bone; o, occipital bone. Scale bars, 5 mm.

To investigate postnatal changes in cranial sutures over time, skulls of *Runx2^+/+^* and *Runx2^m/+^* mice were analyzed at 1, 2, and 3 months of age using μCT, and 3D rendered images were examined (Fig. 6E). At 1 month, while cranial sutures of *Runx2^+/+^* mice were nearly closed, those in *Runx2^m/+^* mice remained open, showing delayed closure and Wormian bones at the lambdoid suture (Fig. 6E). At 2 and 3 months, the sutures in *Runx2^m/+^* mice remained open with persistent closure defects and Wormian bones, indicating impaired cranial ossification (Fig. 6E). Consistently, skeletal preparations at 6 months showed that *Runx2^m/+^* mice had an open anterior fontanelle, impaired suture closure, a short nasal apex, and zygomatic arch dysplasia, whereas *Runx2^+/+^* mice had fully closed cranial sutures (Fig. 6F).

Taken together, these findings demonstrate that *Runx2^m/+^* mice carrying a clinically relevant RHD missense mutation develop postnatal features of severe CCD.

### Formation of accessory root-like protrusions in *Runx2^m/+^* mice

CCD patients often exhibit delayed eruption and supernumerary teeth^(1)^, whereas supernumerary teeth are absent in heterozygous *Runx2*-deficient mice^(12)^. Similarly, no supernumerary teeth were observed in *Runx2^m/+^* mice; however, 3D rendering of μCT images revealed a previously unreported accessory root-like protrusion between the three roots of the maxillary first molars (Fig. 7A-D).

**Fig. 7.**
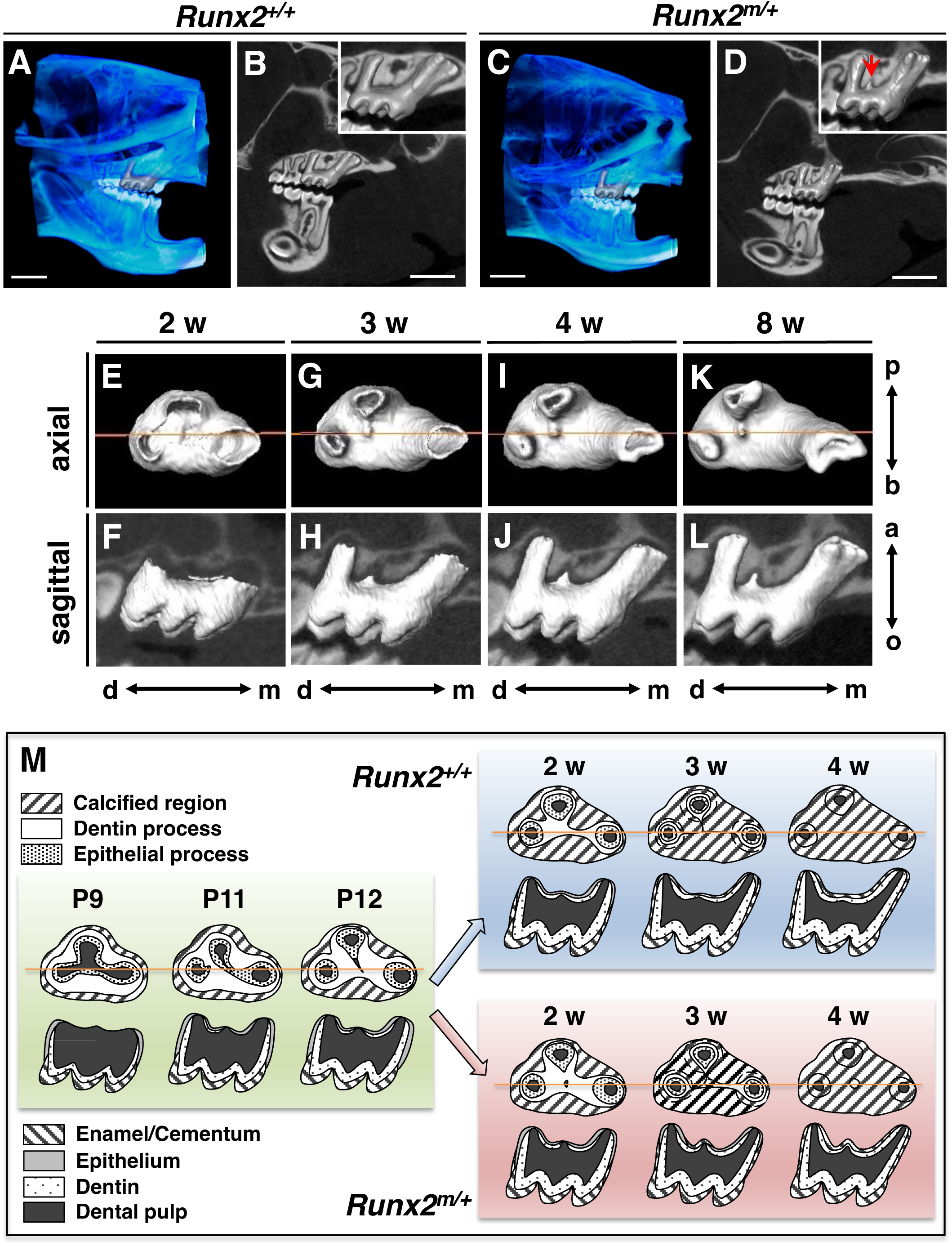
Dental phenotypes of postnatal *Runx2* missense mice. (A-D) 3D µCT images of the maxilla and mandible in *Runx2^+/+^* and *Runx2^m/+^* mice at 3 months. Panels (A) and (C) show whole jaw views. Panels (B) and (D) show slice images with overlayed 3D reconstructions of the maxillary first molar. Insets display enlarged views of the molars. A red arrow in (D) indicates an additional root-like structure. Scale bars, 2 mm. (E-L) 3D µCT images of the right maxillary first molars in *Runx2^m/+^* mice at 2, 3, 4, and 8 weeks. Axial (E, G, I, K) and corresponding sagittal (F, H, J, L) views from the same individuals are shown. Orange lines in axial panels indicate the sagittal section planes. Directional arrows indicate the palatal (p), buccal (b), apical (a), occlusal (o), distal (d) and mesial (m) directions. Images are from the same individuals (n = 3). (M) Schematic illustration of root furcation in *Runx2^+/+^* and *Runx2^m/+^* mice. Horizontal (top) and sagittal (bottom) views are shown, with orange lines in (E-K) indicating the sagittal section planes. Tissue types are presented by pattern-filled squares. The green panel shows the fusion of three epithelial processes (two buccal and one palatal) at P9, P11, and P12. The blue and pink panels depict the progression of root furcation at 2, 3, and 4 weeks in *Runx2^+/+^* and *Runx2^m/+^*mice, respectively.

In mice, the maxillary first molars develop two buccal roots (mesial and distal) and one palatal root, with root development coordinated through the independent growth of each root from the furcation area via Hartwig’s epithelial root sheath (HERS)^(33)^. The furcation process of the mouse maxillary first molar begins with the formation of the distal buccal and palatal root, followed by separation of the mesial buccal root and palatal roots, mediated through the fusion of epithelial tongue-like processes formed by HERS^(33)^. To examine the development of the small root-like protrusion, 3D μCT images of the maxillary first molars of *Runx2^m/+^* mice were analyzed at 2, 3, 4, and 8 weeks (Fig. 7E-I). At 2 weeks, crown formation was complete, but calcification at the root furcation area was incomplete, and three roots were not yet fully separated (Fig. 7E, F). By 4 weeks, roots were clearly separated and elongated rapidly, with a small protrusion forming near the furcation between the distal buccal and palatal roots (Fig. 7G-J). At 8 weeks, root elongation continued, but the protrusion size remained unchanged (Fig. 7 K, L). Figure 7M illustrates the phenotype observed in this study alongside previous findings in wild-type mice^(33–35)^, and shows that in *Runx2^m/+^*mice, the epithelial processes fail to close properly during root furcation, causing a small protrusion.

Hence, these results identify a previously unrecognized dental phenotype associated with *Runx2* haploinsufficiency and provide new insights into the molecular mechanism underlying root furcation.

## Discussion

This study presents the first *in vivo* functional analysis of pathological *Runx2* RHD mutations using novel mouse lines carrying either a missense or a frameshift-induced dysfunctional RHD. Clinically relevant *Runx2^m/+^* mice harboring p.R232Q corresponding human RUNX2 p.R225Q, exhibited skeletal abnormalities characteristic of classic CCD. Notably, this is the first *in vivo* evidence that homozygous RHD mutations alone can completely abolish bone formation, resulting in a skeleton composed entirely of cartilage. Although supernumerary teeth were absent in heterozygous mutants, 3D μCT imaging revealed a novel accessory root-like protrusion in the maxillary first molar during furcation. These findings demonstrate the importance of modeling specific RHD mutations and provide new insights into the skeletal and dental pathology of CCD beyond previous studies of *Runx2*-deficient mice.

Missense substitutions account for nearly half of all mutations reported in the Human Gene Mutation Database^(36)^. Missense mutation p.R225Q is one of the most frequent across diverse populations worldwide^(12,15,28,37,38)^. Arginine (R) residues, such as R225 in human RUNX2 is crucial for protein stability through hydrogen bonds and salt bridges. Substitutions of these residues can disrupt interactions with charged molecules like DNA and other proteins, leading to pathogenic effects^(39)^. *In silico* analysis predicts that R225, a highly conserved residue across species, forms two hydrogen bonds with E226, which are lost in mutant RUNX2^(38)^. Consistently, the equivalent mouse mutation p.R232Q abolishes osteocalcin promoter activation due to loss of RHD function. In homozygous mice, expression of *Col10a1*^(22)^, a direct Runx2 target, was markedly reduced. The p.R232Q Runx2 failed to sustain *Runx2* expression in hypertrophic chondrocytes, causing maturational arrest. In *Runx2^Δ2/Δ2^* mice, the lack of bone formation results from loss of function, most likely due to nonsense-mediated decay triggered by the frameshift mutation, consistent with previous reports that nonsense or frameshift mutations also cause classic CCD^(5,40,41)^.

Two *Runx2* knockout mouse lines have been previously reported: one generated by disrupting the Q/A repeat domain with an *IRES-lacZ-pA* cassette^(12)^, and another by deleting an exon containing 41 amino acids of the RHD using a *PGK-neo* cassette^(7)^. In the latter line carrying a LacZ reporter, X-gal staining showed persistent activity of the endogenous *Runx2* promoter even in homozygotes, with blue staining in bone-forming regions^(12)^. However, the prolonged half-life of LacZ hampers accurate tracking of the temporal dynamics of mutant Runx2 expression during skeletogenesis. Indeed, *Runx2^m/m^*embryos exhibited markedly reduced mutant Runx2 expression, suggesting that loss of functional Runx2 impairs either the maintenance of its transcription or protein stability. These findings shed light on the *in vivo* regulation and dynamics of Runx2 expression that are not captured by reporter-based approaches.

The classical NLS typically comprises 5-20 amino acids rich in basic amino acid residues like lysine (K) and arginine (R)^(42)^. Runx2 contains a Myc-related 9 amino acid NLS (PRRHRQKLD) at the RHD-PST junction, essential for its nuclear localization^(32)^. Previous studies using NIH-3T3, U2OS, C3H10T1/2, and HEK293T cells showed that wild-type RUNX2 localizes to the nucleus, whereas p.R225Q or p.R225W mutants exhibited impaired nuclear translocation^(37,43)^. Fusion proteins carrying p.R225Q or p.R225W mutation fail to efficiently accumulate in the nucleus^(37)^, unlike the wild-type. RUNX2 carrying either the p.K218N or p.K220I mutation (previously described as K204N and T206I, respectively), both located near the NLS, fails to translocate to the nucleus. In contrast, the p.R225Q mutant (previously described as R211Q) shows approximately equal distribution between the nucleus and cytoplasm in about half of the cells^(43)^. Although R232 in mouse Runx2 is located near the NLS, its mutation caused only a partial defect in nuclear localization, with approximately 70% of the mutant Runx2 detected in the nuclei of hypertrophic chondrocytes in *Runx2^m/m^* mice. Thus, reduced function and expression due to RHD dysfunction, rather than impaired nuclear localization, is likely the primary contributor to the skeletal phenotype. A missense mutation in the RHD has been reported to exert a dominant-negative effect^(44)^, which may explain the observation that the mineralized area is smaller in missense mutant compared to the frameshift mutant.

In CCD patients, supernumerary teeth most commonly develop in the incisors and premolars^(45)^. In contrast, mice have a monomorphic dentition with one continuously growing incisor and three molars, separated by a toothless diastema, lacking canines or premolars^(46)^. Unlike the dimorphic dentition of humans, tooth eruption in *Runx2*-deficient mice is only slightly delayed or not significantly affected^(9,47)^, similar to the normal eruption of deciduous teeth in CCD patients^(2,48)^. These species-specific differences in dental architecture may explain why the dental phenotype in mice is less pronounced than the skeletal phenotype.

We identified an accessory root-like protrusion in the maxillary first molars of *Runx2^m/+^* mice, representing a novel dental phenotype distinct from previously reported abnormalities such as short roots or taurodontism^(49)^. In normal maxillary molars, horizontal differential growth of HERS forms three tongue-like interradicular processes that fuse to divide the single cervical opening into three parts, initiating multi-root formation^(33,50)^. The accessory roots likely result from impaired fusion of these processes at the leading edge of HERS, caused by *Runx2* haploinsufficiency (Fig. 7M). As Runx2 is critical for epithelial-mesenchymal interactions during root development, reduced expression may disrupt spatiotemporal regulation of HERS fusion during furcation, leading to protrusion formation. Notably, this protrusion was much smaller than normal roots, suggesting incomplete root elongation, possibly due to aberrant odontoblast differentiation. Given that maxillary first molars normally have three roots and the most complex furcation among molars, their intricate morphology and spongy maxillary bone structure may underlie susceptibility to furcation abnormalities in *Runx2^m/+^* mice.

In conclusion, our *Runx2* mutant mice replicate key skeletal features of CCD and uncover a novel dental phenotype characterized by accessory roots, likely resulting from impaired root furcation. Although unconfirmed in humans, homozygous mutants in mice exhibited a complete lack of bone formation. These findings highlight the critical role of the RHD in both skeletal and multi-root development, providing new insights into CCD pathogenesis.

## Data availability

All data associated with this study are available upon request.

## Acknowledgements

Plasmids for mouse *Col10a1* and *Spp1* were generously provided by Dr. Bjorn R Olsen (Harvard Medical School) and Dr. Shinsuke Ohba (Osaka University), respectively. The *p6OSE2-luc* vector was obtained from Dr. Karsenty (Columbia University) through Dr. Kenji Hata (Osaka University). We thank Drs. Kosuke Hosoba and Xynyi Yu for their technical assistance. Parts of this work was performed using the facilities at the Natural Science Center for Basic Research and Development and the Central Laboratory, Faculty of Dentistry, Hiroshima University. This study was supported by the Japan Society for the Promotion of Science Grants-in-Aid for Scientific Research (Grant Numbers: 16K15780, 21KK0161, 22K10148, 25K13145); the Frontier Development Program for Genome Editing funded by the Doctoral Program for Worldleading Innovative and Smart Education, JST SPRING (Grant Number JPMJSP2132); and the Cooperative Research Program of the Institute for Frontier Life and Medical Sciences, Kyoto University, Japan.

## Author contributions

C.S. designed the research; S.Ogawa, S.H., Y.Y., M.H., C.S., A.H., H.W., T.S., K.H., S.Ogata, K.F., and G.K. performed the experiments; C.S., S.Ogawa, S.H., Y.Y., A.H., S.M., and T.S. analyzed data; C.S., S.Ogawa, S.H., and Y.Y. wrote the paper; C.S., S.M., K.U., T.O., K.T., T.K., and D.D. provided feedback and guidance; S.M., A.H., T.S., K.F., T.O., T.Y., R.K., Y.S., and D.D. reviewed and edited the manuscript.

