## Supplementary Material for "Functional impact of pathogenic Runt domain mutations in *Runx2* on skeletal and dental development in cleidocranial dysplasia"

Supplementary figures S1 to S7 with legends and Supplementary Tables S1-S3

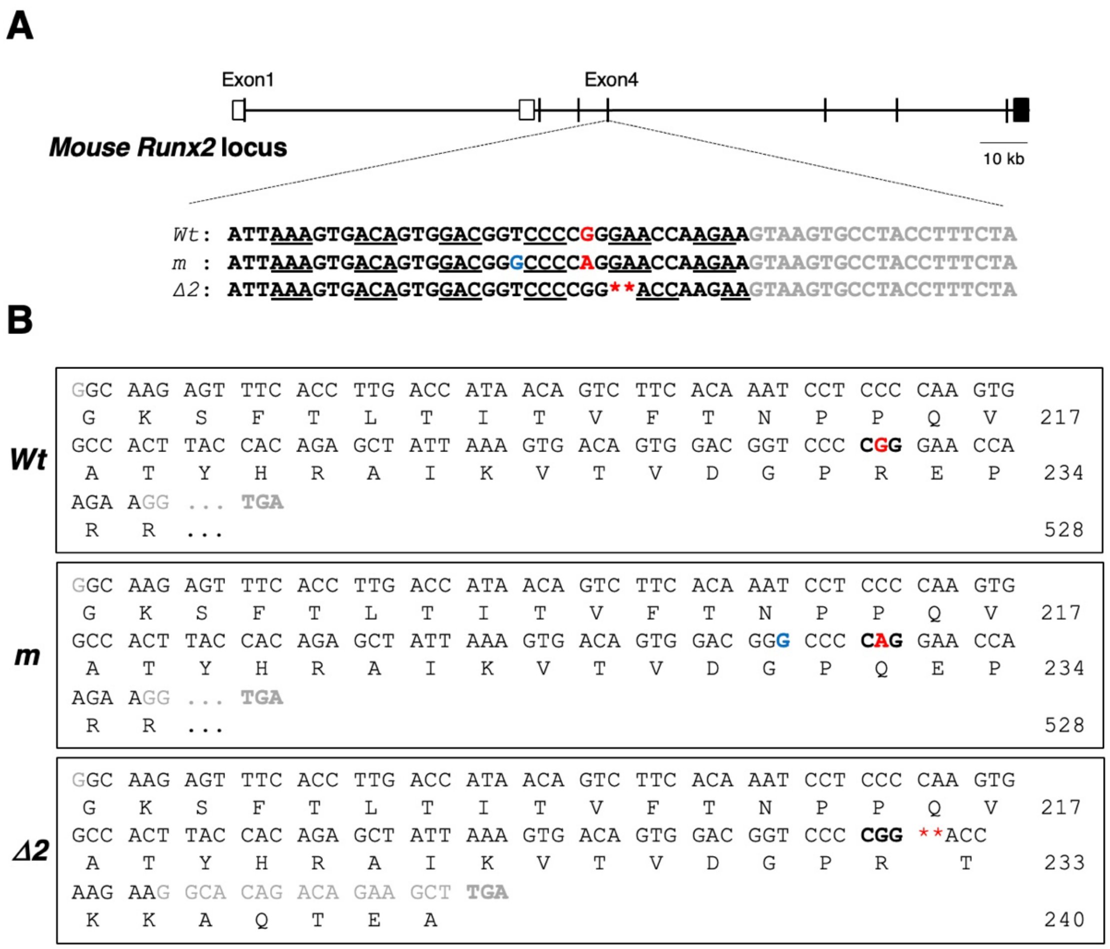


**Supplementary figure 1.** Amino acid sequences and mutation sites in three established *Runx2* mutant mice. (a) Sequences of the wild-type of *Runx2* allele (*Wt*) and two mutant alleles (*m* and *Δ2*) introduced into exon 4 of the mouse *Runx2* gene. (b) The amino acid sequences of each mutant compared to the wild-type mouse *Runx2*. In the *m* allele, a missense mutation changes arginine (R) at position 232 to glutamine (Q), a silent mutation changes the nucleotide at position 230 from thymine (T) to guanine (G), but the encoded amino acid remains glycine (G). The amino acid at position 232 is shown in bold. The nucleotide with the silent mutation at position 230 and the missense mutation at position 232 are shown in bold black and bold red, respectively. (c) In the *Δ2* allele, deletion of the first two nucleotides of the codon encoding glutamate (E) at position 233 causes a frameshift, resulting in a premature stop codon (TGA) at position 241. Nucleotide deletions are indicated by asterisks.

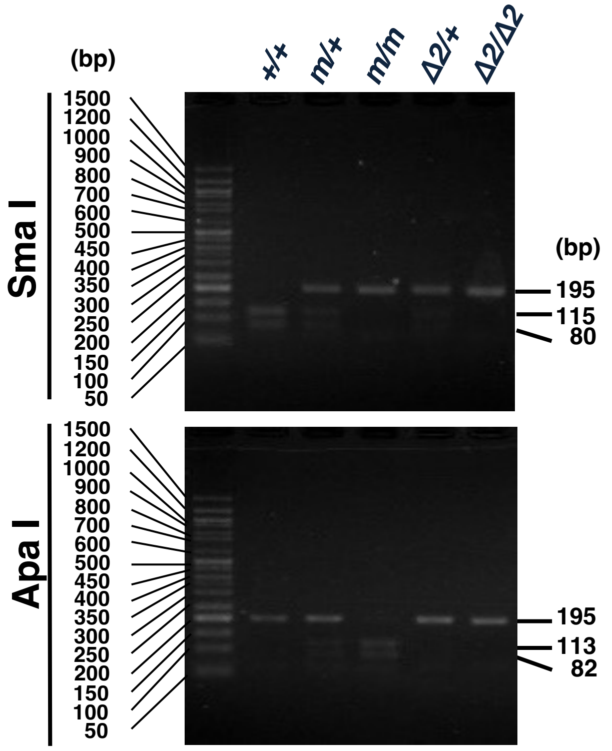


**Supplementary figure 2.** Genotyping of wild-type and *Runx2* mutant mice by PCR and restriction enzyme digestion. Genotyping PCR for wild-type and mutant mice is shown. DNA extracted from ear punches was amplified using primers described in the Materials and Methods. The 195 bp product amplified from the *Wt* allele is digested with SmaI into 115 bp and 80 bp fragment, whereas the product amplified from the mutant alleles (*m* and *Δ2*) remain undigested. The PCR product from the m allele is digested with Apa I into 113 bp and 82 bp fragments. The 195 bp products from the *Wt* and *Δ2* alleles remain undigested by ApaI. A molecular size marker is shown in the leftmost lane.

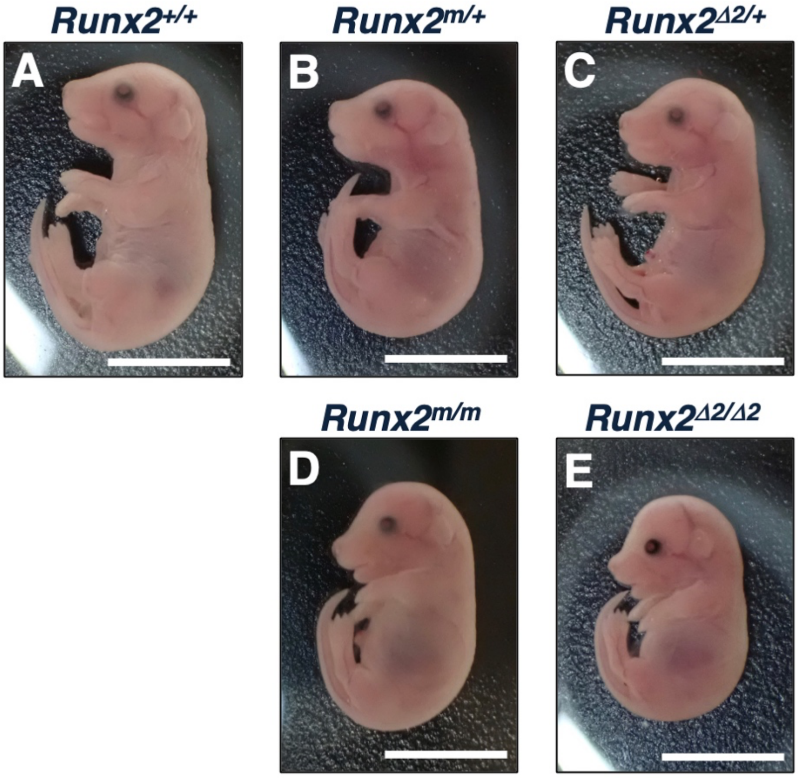


**Supplementary figure 3.** Gross appearance of E18.5 mouse embryos with wild-type and *Runx2* mutant alleles. Representative whole-body images of *Runx2*$^{+/+}$ (a), *Runx2*$^{m/+}$ (b), *Runx2*$^{\Delta2/+}$ (c), *Runx2*$^{m/m}$ (d) and *Runx2*$^{\Delta2/\Delta2}$ (e) embryos at E18.5 are shown. A representative image selected from > 3 neonates per genotype is shown. Scale bars, 10 mm.

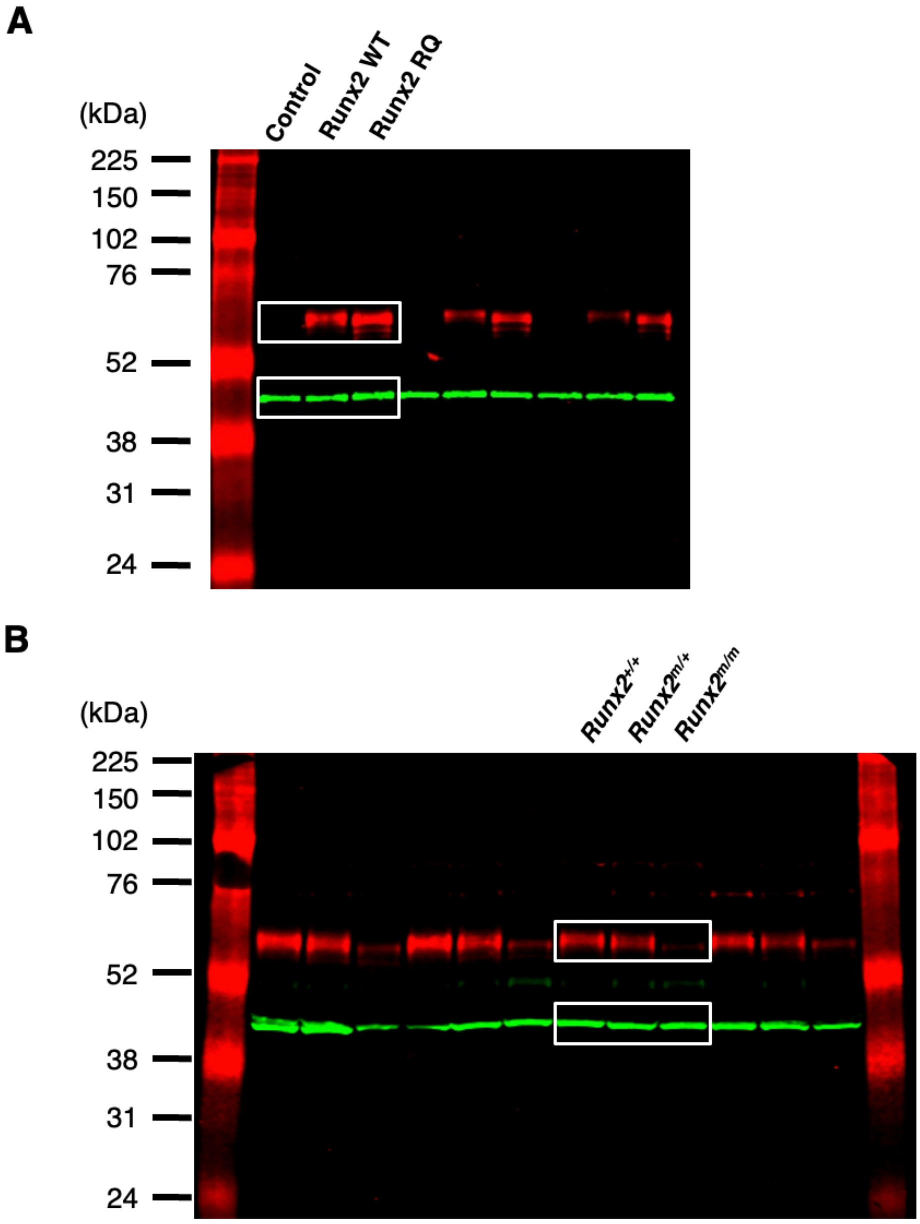


**Supplementary figure 4.** Full-length fluorescent Western blot images of Runx2 and β-actin. Overlay of the full-length Western blot images showing fluorescent signals for Runx2 (700 nm channel, red) and b-actin (800 nm channel, green) of HEK 293T cells (a) and mouse forelimbs (b). Cropped regions are indicated by while boxes, and the processed images are presented in Figs. 4a and 4b.

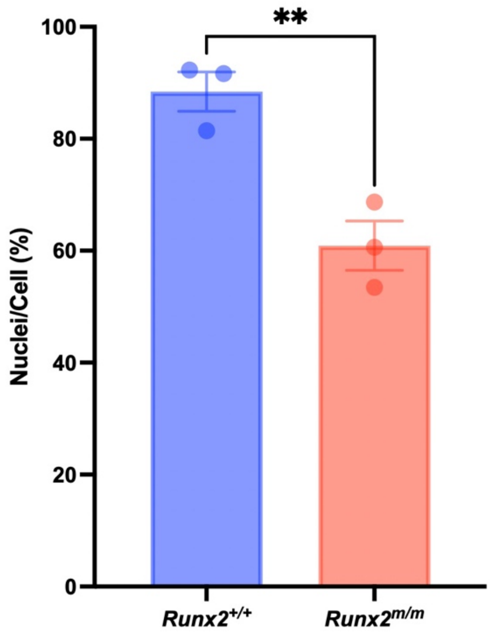


**Supplementary figure 5.** Nuclear translocation of Runx2 in wild-type and *Runx2* mutant embryos. Quantification of the percentage of Runx2 signal localized to the nucleus in *Runx2*$^{+/+}$ and *Runx2*$^{m/m}$ embryos at E18.5. Each bar represents the average from three independent immunohistochemical analyses (three different mice/genotype), with individual values shown as dots (means ± SE). **P < 0.01 vs. *Runx2*$^{+/+}$.

| Injected concentration (mRNA/ssODN, ng/µL) | No. of newborns | NHEJ events (%) | Insertion (%) | Deletion (%) | HDR with ssODN (%) |
|---|---|---|---|---|---|
| 1 / 15* | 18 | 5 (27.8) | 0 (0) | 5 (27.8) | 0 (0) |
| 2 / 15* | 12 | 3 (25.0) | 1 (8.3) | 2 (16.7) | 0 (0) |
| 4 / 100* | 21 | 6 (28.6) | 0 (0) | 6 (28.6) | 2 (9.5) |
| Total | 51 | 14 | 1 | 13 | 2 |
| Established Lines | 5 | **1** | - | 2 | **1** |

**Supplementary table 1.** Genome editing outcomes following mRNA/ssODN microinjection.

*Concentrations of mRNA and ssODN used during microinjection. NHEJ: Non-homologous end joining; HDR: Homology-directed repair using ssODN. Percentages indicate the proportion of newborns with the respective genome editing outcome.

| Nucleotide Change | Mutation Type | Amino Acid Change Mouse/ Human | Primer Sequences (5’→3’) |
|---|---|---|---|
| None | Wild-type | No change | F: ATGGCGTCAAACAGCCTCTTCAGCG<br>R: TCAATATGGCCGCCAAACAGACTCATCC |
| G→A | Missense | p.R232Q /p.R225Q | F: TCCCCAGGAACCAAGAAGGCACAGACAGAAGC<br>R: CCGTCCACTGTCACTTTAATAGCTCTGTGGTAAG |
| C→T | Missense | p.R232W/p.R225W | F: TCCCTGGGAACCAAGAAGGCACAGACAGAAGC<br>R: CCGTCCACTGTCACTTTAATAGCTCTGTGGTAAG |
| C→T | Nonsense | p.R200X /p.R193X | F: CGGATGAGGCAAGAGTTTCACCTTGACCATAAC<br>R: CTCCGGCCCACAAATCTCAGATCGTTGAAC |

**Supplementary table 2.** Primer sequences for introducing point mutations into the mouse *Runx2* gene by site-directed mutagenesis. Primers listed in this table were designed for site-directed mutagenesis to introduce specific nucleotide substitutions in the mouse *Runx2* gene. Each mutation is located in the following exons: p.R232Q and p.R232W in exon 4, and p.R200X in exon 3. The Amino Acid Change column shows the resulting substitution in both mouse and human Runx2 proteins. Forward (F) and reverse (R) primers are given in 5' to 3' orientation.

| Gene | Primer Sequences (5’→3’) |
|---|---|
| *mRunx2* | F1: GTTTCCTCTCTTAAATCTGCAGGC<br>R1: CAGAGTGAATTCTTCACAGGGTCC |
| *mCol2a1* | F1: GGACGCTACACTCAAGTCACTG<br>R1: GCGTCTGACTCACACCAGATAG |
| *mIhh* | F1: GCGGACAATCATACAGAACCAG<br>R1: GGGCAGCCTCTCTTCATACTGT |

**Supplementary table 3.** Primer sequences for genotyping and amplification of cDNA fragments. Primers F1 and R1 for mouse *Runx2* (*mRunx2*) were used for genotyping. Primers F1 and R1 for mouse *Col2a1* (*mCol2a1*) and *Indian hedgehog* (*mIhh*) were used to generate templates for RNA probe synthesis by *in vitro* transcription for *in situ* hybridization.
